# Mitophagy contributes to alpha-tocopheryl succinate toxicity in GSNOR-deficient hepatocellular carcinoma

**DOI:** 10.1101/867846

**Authors:** Salvatore Rizza, Luca Di Leo, Sara Mandatori, Daniela De Zio, Giuseppe Filomeni

## Abstract

The denitrosylating enzyme *S-*nitrosoglutathione reductase (GSNOR), has been reported to control the selective degradation of mitochondria through mitophagy, by modulating the extent of nitric oxide-modified proteins (*S-*nitrosylation). The accumulation of *S-*nitrosylated proteins due to GSNOR downregulation is a feature of hepatocellular carcinoma, causing mitochondrial defects that sensitize these tumors to mitochondrial toxins, in particular to mitochondrial complex II inhibitor alpha-tocopheryl succinate (αTOS). However, it is not known if mitophagy defects contribute to GSNOR-deficient cancer cells sensitivity to αTOS, nor if mitophagy inhibition could be used as a common mechanism to sensitize liver cancers to this toxin. Here, we provide evidence that GSNOR-deficient cancer cells show defective mitophagy. Furthermore, we show that αTOS is a mitophagy inducer and that mitophagy defects of GSNOR-deficient liver cancer cells contribute to its toxicity. We finally prove that the inhibition of mitophagy by depletion of Parkin, a pivotal ubiquitin ligase targeting mitochondria for degradation, enhances αTOS toxicity, thus suggesting that this drug could be effective in treating mitophagy-defective tumors.

## 1. Introduction

Mitophagy, the selective targeting of damaged mitochondria for lysosomal degradation, is an essential process controlling mitochondrial homeostasis. Being at the crossroads of metabolic, redox and cell death pathways, mitochondria play a central role in physiology and, not surprisingly, in several pathological settings, including cancer [1–3]. Although dysfunctional mitochondria are a feature of many malignancies, and contribute to the increased glycolytic metabolism referred as “Warburg effect” [4–6], it is becoming clear that these organelles may also contribute to tumorigenesis by i) producing signaling molecules [7–9]; ii) controlling Ca^2+^ homeostasis [10]; iii) acting as hub of cell death signals [11,12]; iv) catabolizing lipids and amino acids for energy production, or v) stimulating their *de novo* synthesis to generate biomass in proliferating cells [13,14]. Due to these multiple roles, changes in the mitochondrial mass, which mostly derives from an imbalance between biogenesis and mitophagy, has been linked to both tumorigenesis and cell survival, as well as to tumor suppression [15]. Although mitophagy appears to be defective during cancer initiation, it acts mainly as a cytoprotective mechanism during cancer progression, as it supports cell survival under stressful conditions, *e.g.* nutrient deprivation or hypoxia [1]. From a clinical point of view, such a cytoprotective role can contribute to chemoresistance [16–18]. In support to this, BCL2-interacting protein 3 like (NIX)-dependent mitophagy has been identified as a process attenuating doxorubicin toxicity in colorectal cancer [19]. Similarly, the upregulation of ariadne RBR E3 ubiquitin protein ligase 1 (ARIH1) has been demonstrated to protect lung cancers from cisplatin by interacting with the mitochondrial potential sensor PTEN-induced kinase 1 (PINK1), and stimulating a Parkin-independent mitophagy [16].

Several studies tested mitophagy inhibition as a method to enhance drugs sensitivity [20,21], and it has been suggested that selective silencing of mitophagy genes (*e.g.* PINK1 [16], Parkin [22,23], BCL2/adenovirus E1B 19 kDa protein-interacting protein 3, BNIP3 [24,25]), or the inhibition of lysosomes function (by chloroquine [26], or the alkaloid liensinine [27]) potentiates the antitumoral activity of conventional chemotherapeutics, *e.g.* doxorubicin [27], salinomycin [28], paclitaxel and vincristine [29]. Notwithstanding this amount of data, to date, the use of specific mitophagy inhibitors is still far from being implemented in combined anticancer therapies.

We have recently demonstrated that the denitrosylase *S-*nitrosoglutathione reductase (GSNOR) indirectly regulates a set of proteins – such as Parkin [30,31] – that are post-translationally modified by nitric oxide (NO) through a reaction called *S-*nitrosylation [32,33]. Parkin *S-*nitrosylation inhibits its capability to target damaged mitochondria for mitophagy degradation [34,35]. As a result, cells in which GSNOR is downregulated are mitophagy-defective [30].

GSNOR is a highly conserved enzyme that has been found downregulated in human hepatocellular carcinomas [36,37] and breast cancer [38]. The etiological role of GSNOR in tumorigenesis is also confirmed by the evidence that GSNOR knock-out (KO) mice spontaneously develop liver cancer [36]. However, if this depends on mitophagy defects has not been investigated yet. In a previous work, we also demonstrated that GSNOR-KO cells show mitochondrial defects that can be exploited to selectively kill hepatocellular carcinoma cells [39] by the use of a class of mitochondrial drugs (named mitocans [40]) directed to the Complex II of the mitochondrial respiratory chain (also known as succinate dehydrogenase, SDH).

Here we provide evidence that mitophagy is involved in the sensitivity of GSNOR-KO liver cancer cells to mitochondrial toxins, *i.e.* alpha-tocopheryl succinate (αTOS) [41–43], suggesting the use of SDH-directed mitocans for the treatment of mitophagy-defective tumors.

## 2. Materials and Methods

### 2.1. Cell culture and treatments

HepG2 cell line was obtained from the Banca Biologica e Cell Factory (IRCCS AOU San Martino - IST Istituto Nazionale per la Ricerca sul Cancro). Huh-7 12 (HUH7) were purchased from Sigma-Aldrich. Both cell lines were grown in RPMI 1640 supplemented with 10% FBS and antibiotics at 37°C in an atmosphere of 5% CO_2_. All cell lines were cultured for fewer than 2 months after resuscitation and were used from the third to the fifteenth passage in culture. Cell lines validation was carried out by the producer by means of DNA Profile STR (Short Tandem Repeat) and mycoplasma contamination was routinely screened by a PCR-based assay. All the cell culture media and supplements were purchased from Gibco, Thermo-Fisher Scientific.

Compounds and concentrations used in the study are as follows: carbonyl cyanide m-chlorophenyl hydrazone (CCCP, Sigma-Aldrich), 2.5-10 μM; chloroquine (CQ, Sigma-Aldrich), 20 μM; alpha-tocopheryl succinate (αTOS, Sigma-Aldrich), 40-60 μM. Incubation times are indicated in the figure legends.

### 2.2. Gene silencing

Transient knock down was performed by small interference RNA (siRNA) technique. Cell were transfected using RNAiMAX (Thermo Fisher Scientific) using endonuclease-prepared pools of siRNAs (esiRNA, Sigma-Aldrich) directed against GSNOR (siGSNOR) and Parkin (siParkin), or, as control, with a scramble siRNA duplex (siScr). HepG2 cells stably expressing eGFP-short-hairpin RNAs (shRNA) were generated in our lab with a procedure reported in [39].

### 2.3. Analysis of cell viability and cell death

Dead cells were evaluated by Celigo Imaging Cytometer (Nexcelom Bioscience) upon staining with LIVE/DEAD^®^ Cell Imaging Kit (488/570) (Thermo-Fisher Scientific). The percentage of dead cells was calculated as ratio between propidium iodide-stained cells (dead, red) and the sum of dead cells and calcein AM-positive cells (alive, green).

Cell viability was quantified by reading the fluorescence emission at 590 nm after 2 h incubation with AlamarBlue^®^ Reagent (Thermo-Fisher Scientific) with a Victor X4 (PerkinElmer) plate reader.

Cell survival was assayed upon Cristal Violet staining. Briefly, cells were treated with 40 μM αTOS for 24 h; then they were trypsinized and 1/5 re-seeded in 6-well plates in the presence or not of 40 μM αTOS for 4 days. At the end of the treatment cells were washed twice with PBS, fixed and stained with a solution of 20% (v/v) Methanol (WVR) and 0.05% (w/v) Crystal violet (Sigma-Aldrich) on ice for 10 minutes. After washing, the plate was let completely dry at room temperature. Pictures were acquired with an Olympus microscope equipped with a 4X objective. For quantitation, Crystal violet was eluted with 100% methanol and absorbance measured at 595 nm by a Victor X4 plate reader.

### 2.4. *Analyses of mitochondrial mass and mitochondrial transmembrane potential (*Ψ_*m*_)

Total mitochondrial mass and mitochondrial transmembrane potential (ΔΨ_m_) were analyzed by incubating cells with 50 nM MitoTracker Green-FM or 200 nM TMRM (Thermo-Fisher Scientific), respectively, for 30 minutes in serum-free DMEM. Stained cells were washed twice with cold PBS, collected and analyzed by flow cytometry (FACS Verse, BD-biosciences). Normalized ΔΨ_m_ was calculated as TMRM/MitoTracker Green-FM relative fluorescence (geometric mean).

### 2.5. Immunofluorescence microscopy and analyses

Cells were grown on cover slips preventively coated with Gelatin 1% in PBS (Sigma Aldrich), treated as indicated, washed twice in cold PBS and fixed with 4% paraformaldehyde (VWR) in PBS for 10 min at room temperature.

For mitochondrial dynamics analysis, cells were incubated with a permeabilization solution (PBS/Triton X-100 0.4% v/v), blocked for 1 h with a blocking solution (PBS/normal goat serum 10% v/v) and then incubated for 1 h with and anti-mitochondrial import receptor subunit TOM20 homolog (anti-TOM20 - sc-11415, Santa Cruz Biotechnology) antibody, or, alternatively, with an anti-heat shock protein A9 (anti-GRP75 - ADI-SPS-826, Enzo Life Sciences).

For the evaluation of mitophagy, cells were permeabilized in ice cold methanol (VWR) for 5 minutes, washed three times in PBS, incubated for 1 h in blocking solution and then over night with an anti-Microtubule-associated proteins 1A/1B light chain 3B (anti-LC3, 0231-100, Nanotools) and anti-TOM20 (Santa Cruz Biotechnology) antibodies. Cells were then washed twice with PBS and incubated for 1 h with fluorophore-conjugated secondary antibodies (AlexaFluor 488 and 568). Nuclei were stained with 1□μg/ml Hoechst 33342 (Thermo-Fisher Scientific). Epifluorescence analysis was performed by using Delta Vision (Applied Precision) Olympus IX70 microscope. Confocal microscopy experiments were performed by using LSM800 microscope (ZEISS) equipped with an oil-immersion 63X objective and ZEN imaging software. Fluorescence images were adjusted for brightness, contrast and color balance by using Fiji [44] analysis software. Confocal microscopy images were deconvoluted using the software Huygens Professional (Scientific Volume Imaging).

3D-rendering of multi-stacks images was achieved by UCSF CHIMERA (Reagents of the University of California) [45]. Mitophagy rate was assessed upon incubation with 20 mM chloroquine, added 2 h before the end of the treatments to block mitochondria degradation within the autophagosomes. The percentage of mitochondria particles colocalizing with LC3 puncta was calculated by Fiji analysis software using the open-source plugin ComDet v. 0.3.7. on at least 8 different cells/experimental condition. For both TOM20 and LC3 fluorescence channels, the parameters utilized were: Particle size ≥ 4 px; intensity threshold = 3. The colocalization was considered positive if the maximum distance between the center of 2 particles was ≤ 4 px. The pictures showed represent 3D-projections and were obtained by summing the fluorescence signal of the central z-Stacks (3 planes, 0.3 μm).

### 2.6. Live-imaging confocal microscopy

Live-imaging monitoring of mitophagy was achieved by growing the cells on 96-well flat-bottom plates for microscopy (Greiner). Cells were treated with 40 μM αTOS where indicated for 24 h. Cells were then stained for 30 minutes with 50 nM MitoTracker Green-FM, 200 nM LysoTracker Red (Thermo-Fisher Scientific) and 50 nM Nuclear Violet (Biomol). Then they were washes in PBS, replaced in cell medium and treated with 5 μM CCCP or 40 μM αTOS and placed in the incubator chamber of the ImageXpress Micro Confocal High-Content Imaging System (Molecular Devices) at 37°C in an atmosphere of 5% CO_2_. Pictures were captured every 5 minutes for 60-90 minutes.

### 2.7. Transmission electron microscopy

HepG2 cells stably expressing an eGFP-short-hairpin against GSNOR or a scramble non-target sequence, were treated with 40 μM αTOS for 24 h, then fixed with 2.5% glutaraldehyde (Sigma-Aldrich) buffered with 0.1 M sodium phosphate, pH 7.4. Sections were prepared at the Core Facility for Integrated Microscopy (CFIM), Copenhagen University and images were acquired with a CM 100 BioTWIN electron microscope equipped with an Olympus Veleta digital camera. Figures were processed with an Olympus ITEM software.

### 2.8. Protein determination

Protein concentration was determined by the method of Lowry [46].

### 2.9. Western blot

Samples preparation and acquisition of Western blots were performed as previous reported [39]. Western blots shown are representative of at least n=3 independent experiments giving similar results. We used the following primary antibodies: anti-succinate dehydrogenase subunit A (anti-SDHA - ab14715, 1:5000) from Abcam; anti-TOM20 (sc-11415, 1:5000), anti-nitric oxide synthase 2 (anti-NOS2 - sc-8310 1:1000), anti-GSNOR (sc-293460, 1:1000), anti-Parkin (sc-30130, 1:1000), anti-Voltage-dependent anion channel (anti-VDAC - sc-8828, 1:2000) from Santa Cruz Biotechnology; anti-Vinculin (V4505, 1:10000) from Sigma-Aldrich; anti-thioredoxin 1 (anti-Trx1 - 2298, 1:1000) from Cell Signaling.

### 2.10. Statistical analyses

Values are expressed as means ± SD (or SEM where specified) and statistical significance was assessed by Student’s t-test using Prism 8.0 (GraphPad Software, Inc.) in order to determine which groups were significantly different from the others.

## 3. Results

### 3.1. GSNOR-deficient hepatocellular carcinoma cells show defective mitochondrial dynamics

In order to study the effects of GSNOR-ablation on mitochondria in hepatocellular carcinoma, we took advantage of two different hepatoma cell lines: HUH7, in which we transiently knocked-down GSNOR by RNA interference, and HepG2, stably expressing eGFP-shRNAs targeting GSNOR (shGSNOR), or a non-target sequence (shScr) as a control. As shown in **Figure 1A**, neither transient nor stable GSNOR ablation affected the expression levels of nitric oxide synthase 2 (NOS) - the predominant NO-producing enzyme in HUH7 and HepG2 cells - and thioredoxin 1 (Trx1), which might complement lack of GSNOR denitrosylating activity induced by sh or siRNAs. These data suggest that no compensatory mechanisms are induced by GSNOR downregulation in our experimental conditions. However, as previously reported in murine neurons and fibroblasts [30,31], GSNOR knocking-down resulted in a marked decrease of mitochondrial transmembrane potential (Ψ_m_) (**Figure 1B**) and a fragmentation of mitochondrial network (**Figures 1C and D**). Overall, these data suggest that, in hepatocellular carcinoma cells, GSNOR deficiency results in mitochondrial fragmentation and depolarization.

**Figure 1.**
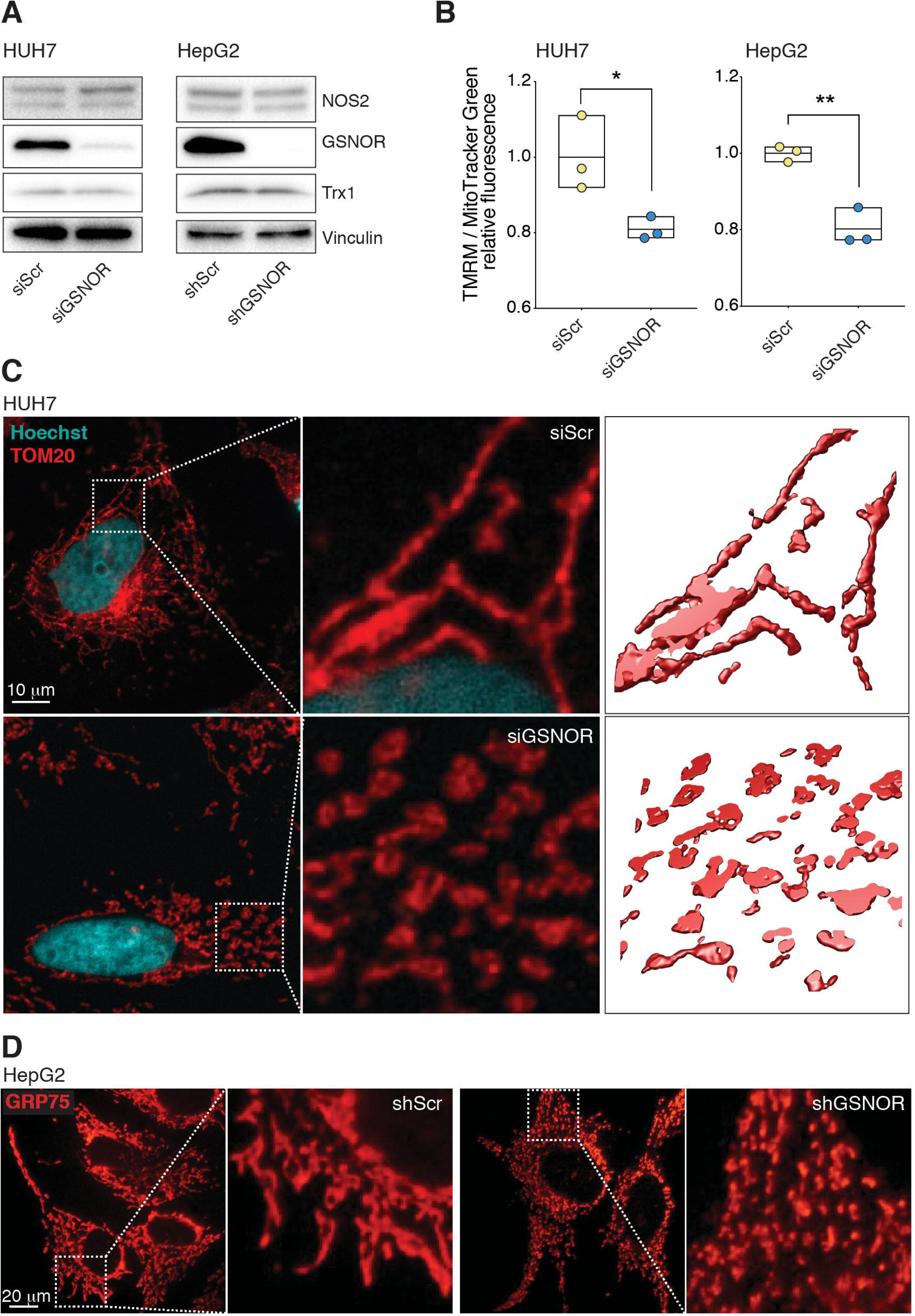
GSNOR-deficient hepatocellular carcinoma cells show defective mitochondria. **(A)** Western blot analysis of NOS2, Trx1 and GSNOR in HUH7 cells transiently downregulating GSNOR (siGSNOR) and HepG2 cells stably expressing eGFP-shRNAs against GSNOR (shGSNOR), along with their relative scrambled RNA-trasfected counterparts (siScr, shScr respectively). Vinculin was selected as loading control. **(B)** Mitochondrial transmembrane potential (ΔΨ_m_) in HUH7 and HepG2 cells evaluated by flow cytometric analysis of TMRM fluorescence. Values are normalized on the total mitochondrial mass (MitoTracker Green relative fluorescence intensity) and expressed as fold change. Values represent the means ± SD of n = 3 independent experiments. \**p* < 0.05; \*\**p* < 0.01. **(C)** 3D reconstruction of mitochondrial network in siScr and siGSNOR HUH7 cells revealed by confocal fluorescence microscopy upon incubation with an antibody against the mitochondrial protein TOM20 (red). 3D-rendering of TOM20 signal is shown on the right panels and represents 4-6 z-stacks (0.3 μm size). Hoechst 33342 (blue) was used to visualize nuclei. Scale bar 10 μm. **(D)** Representative fluorescence microscopy analyses of mitochondrial networks of shScr and shGSNOR HepG2 cells performed upon incubation with an antibody against Grp75. Scale bar 20 μm.

### 3.2. GSNOR-deficient hepatocellular carcinoma cells show defective mitophagy

We next investigated the effects of GSNOR downregulation on mitophagy, since this is a mechanism deeply impaired in condition of nitrosative stress induced by GSNOR decrease [30]. To this end, we treated HUH7 cells with the uncoupling agent CCCP, a well-known mitophagy inducer and evaluated mitochondrial mass upon MitoTracker Green staining as indirect measure of mitophagy. (**Figure 2A**). Cytofluorometric assay indicated that CCCP induced a milder decrement of mitochondrial mass in siGSNOR than the control (siScr) counterpart, suggesting that mitochondrial removal was less efficient. To confirm these set of data, we performed live-imaging confocal microscopy analyses of HUH7 cells stained with Lysoracker Red (to follow lysosomes) and MitoTracker Green (to identify mitochondria). Results shown in **Figure 2B** (**Movies 1-6**) indicate that GSNOR-deficiency induced a defective recognition of mitochondria, strengthening the idea that mitophagy rate was compromised in siGSNOR cells. Accordingly, Western Blot analyses performed on HepG2 cells showed that the levels of three mitochondrial proteins commonly used as an estimation of mitochondrial mass (*i.e.* SDHA, VDAC and TOM20) were only slightly affected by CCCP in GSNOR-deficient conditions (shGSNOR) if compared with those measured in control counterparts (shScr) (**Figure 2C**). Overall, these results suggest that loss of GSNOR impairs mitophagy in hepatocellular carcinoma cells.

**Figure 2.**
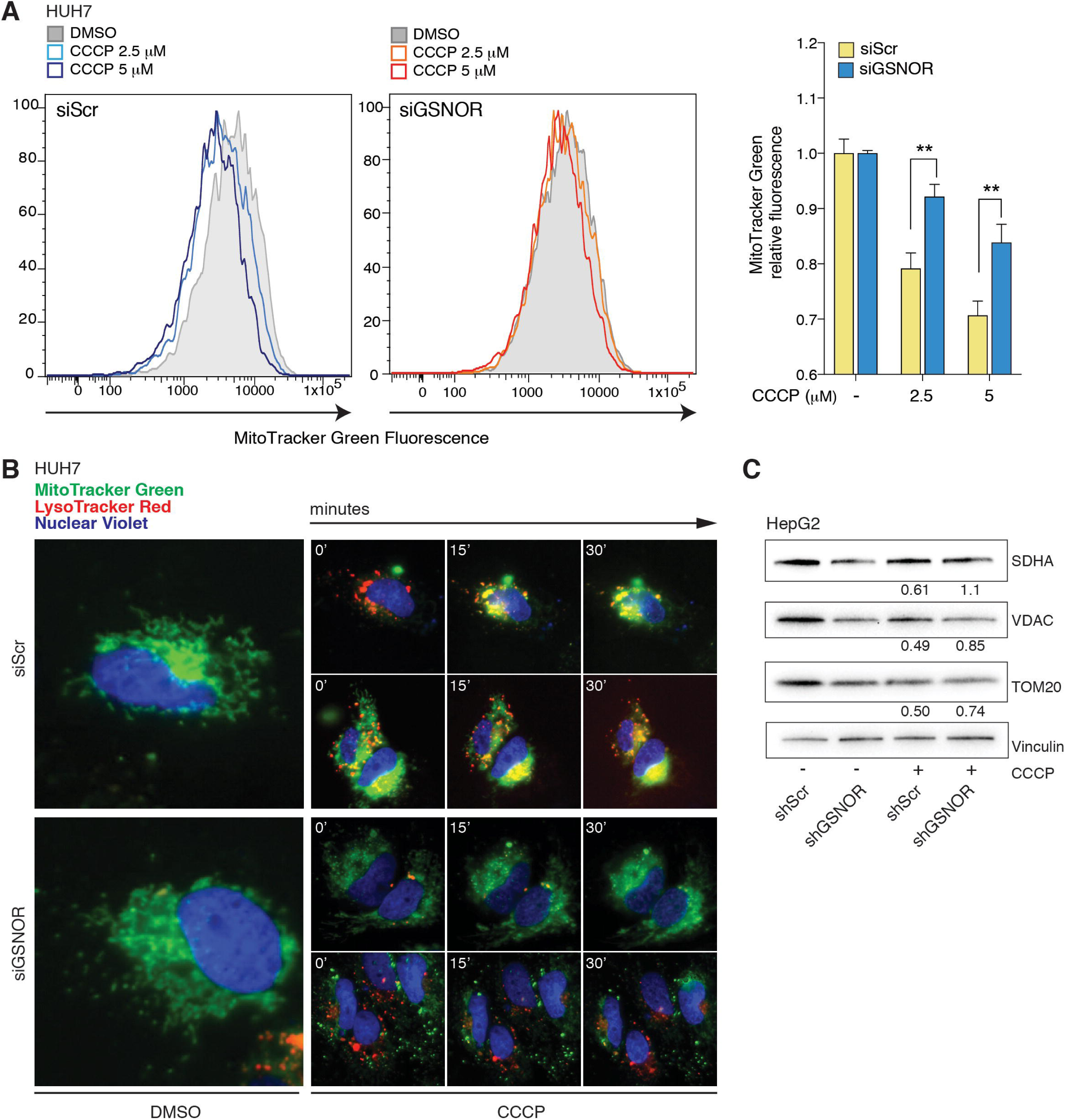
GSNOR-deficient hepatocellular carcinoma cells show defective mitophagy. **(A)** Mitochondrial mass analyzed by flow cytofluorometric detection of MitoTracker Green fluorescence of siScr and siGSNOR HUH7 cells treated for 4 h with 2.5 or 5 μM CCCP. Cytofluorometric histograms are shown on the left. Values (on the right) represent the means ± SEM of n = 3 independent experiments performed in triplicate. \*\**p* < 0.01. **(B)** Representative frames (at 0’, 15’ and 30’) captured upon live-imaging fluorescence microscopy (Movies 1-6) of siScr and siGSNOR HUH7 stained with LysoTracker Red (red), MitoTracker Green (green) and Nuclear Violet (blue) to visualize lysosomes, mitochondria and nuclei, respectively. 5 μM CCCP (or DMSO as vehicle) was added before images acquisition to induce mitophagy. Images were acquired every 5 minutes for 1 h. **(C)** Western blot analysis of succinate dehydrogenase subunit A (SDHA), Voltage-dependent anion channel (VDAC) and Mitochondrial import receptor subunit TOM20 homolog (TOM20) performed on shScr and shGSNOR HepG2 upon treatment with 10 μM CCCP (or DMSO) for 3 h. Vinculin was used as loading control. CCCP/DMSO protein ratio – normalized on Vinculin – is shown below the immuno-reactive bands and indicates the fold decrease of each mitochondrial protein upon CCCP treatment.

### 3.3. GSNOR-deficient hepatocellular carcinoma cells are insensitive to αTOS-induced mitophagy

We previously demonstrated that, in hepatoma cells, GSNOR-deficiency is a condition associated with a rearrangement of the respiratory chain, namely the upregulation of complex II (succinate dehydrogenase, SDH) activity and oxygen consumption [39], presumably as a mechanism to compensate the inhibition of complex I and IV. This feature provided the molecular rationale to the use of SDH-targeting mitochondrial toxins, exemplified by the vitamin E-derivative αTOS that, indeed, showed high (and selective) toxicity towards GSNOR-depleted cells [39].

In the light of the results so far obtained, we wondered if αTOS was able to induce mitophagy and, further, if defects in mitophagy observed in GSNOR-deficient cells could contribute to their enhanced sensitivity towards this drug. To this end, we performed electron microscopy analyses of HepG2 cells treated with αTOS for 24 h and observed that mitochondrial ultrastructure was compromised. In particular, we noticed a decrease in the average mitochondrial size and in the number of mitochondrial cristae (**Figure 3**) in both shScr and shGSNOR cells. Interestingly, in shScr cells we noticed the presence of double-membraned structures surrounding the damaged organelles, presumably early autophagosomes. On the contrary, these structures were absent (or very rare) in GSNOR-deficient cells treated with αTOS, notwithstanding mitochondria shape appeared severely compromised already in control (vehicle-treated) conditions. In order to confirm these results, we performed immunofluorescence analysis by labeling mitochondria with an anti-TOM20 antibody, and autophagosomes with an anti-LC3 antibody. To highlight mitophagy rate, this experiment was performed in the presence of chloroquine (CQ), a well-known inhibitor of lysosome acidification that blocks the degradation of autophagolysosomes content [47]. Fluorescence microscopy analyses indicated that αTOS was able to target mitochondria toward lysosomal degradation in siScr HUH7 cells, but not in GSNOR-deficient cells (**Figure 4A**). Interestingly, the inability to trigger mitophagy was comparable to that observed in cells downregulating Parkin (siParkin) – the ubiquitin-ligase involved in targeting damaged mitochondria for degradation through mitophagy – that were selected as control (**Figure 4A**). An estimation of mitophagy rate that gives the general idea of this phenomenon was provided in **Figure 4B** as unbiased count of the colocalization between anti-TOM20/anti-LC3 double positive particles. In line with these observations, we obtained similar results in HUH7 cells analyzed by live-imaging confocal microscopy, with αTOS treatment triggering mitophagy in siScr but not in siGSNOR cells (**Figure 4C, Movies 7-12**). Flow cytometry analyses were perfomed to quantify the decrease of mitochondrial mass, eventually confirming that siGSNOR cells retained more mitochondria upon αTOS treatment than the siScr counterpart (**Figure 4D**), providing additional evidence that GSNOR-deficient cells were unable to remove αTOS-damaged mitochondria through selective autophagy.

**Figure 3.**
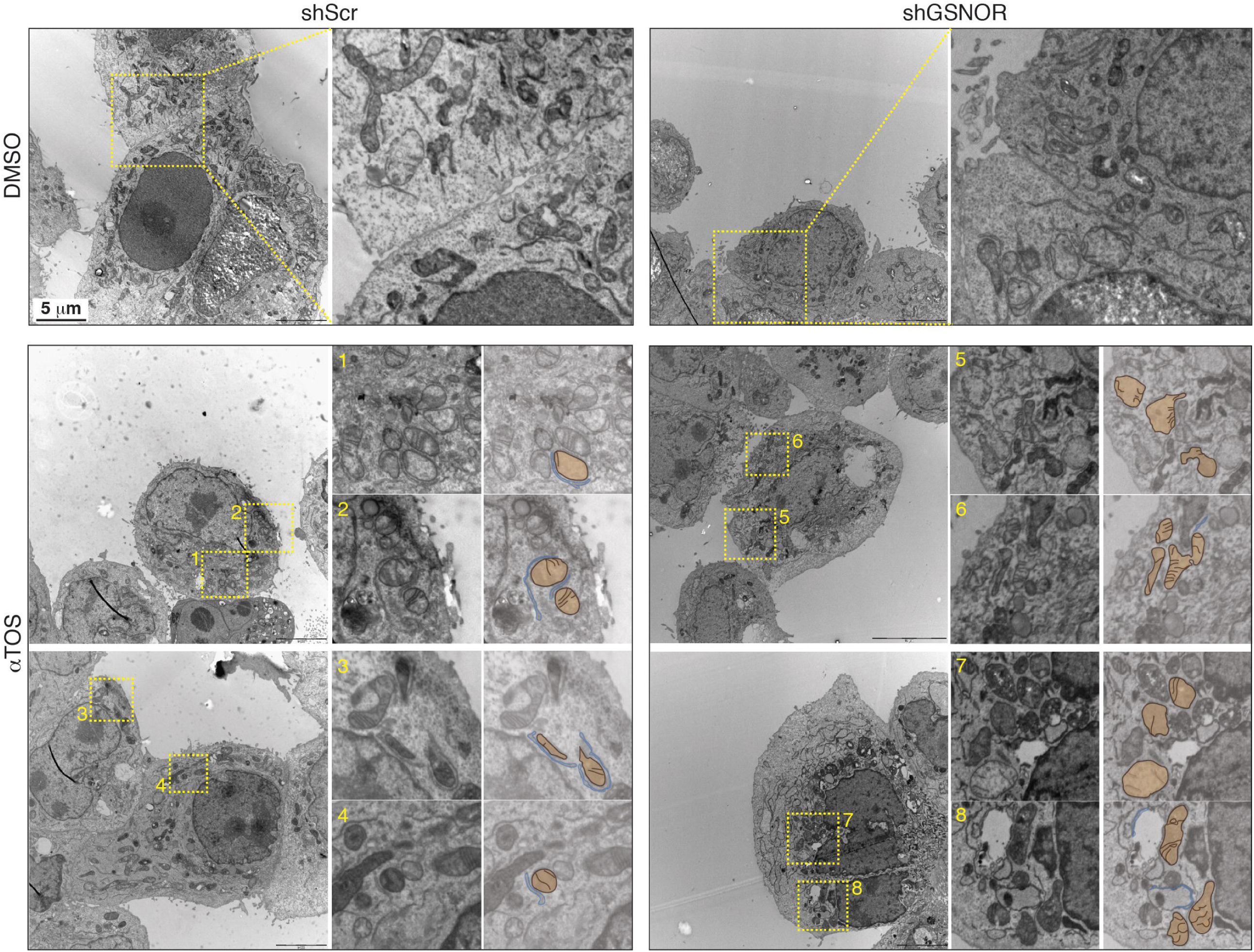
αTOS alters mitochondria structure and induces autophagosome formation. Transmission electron microscopy images of shScr and shGSNOR HepG2 treated for 24 h with 40 μM αTOS or vehicle (DMSO). Magnification of selected fields (yellow dotted squares) are shown on the right panels. Mitochondria are highlighted in light brown and double-membraned structures are highlighted in light blue. Scale bar 5μm

**Figure 4.**
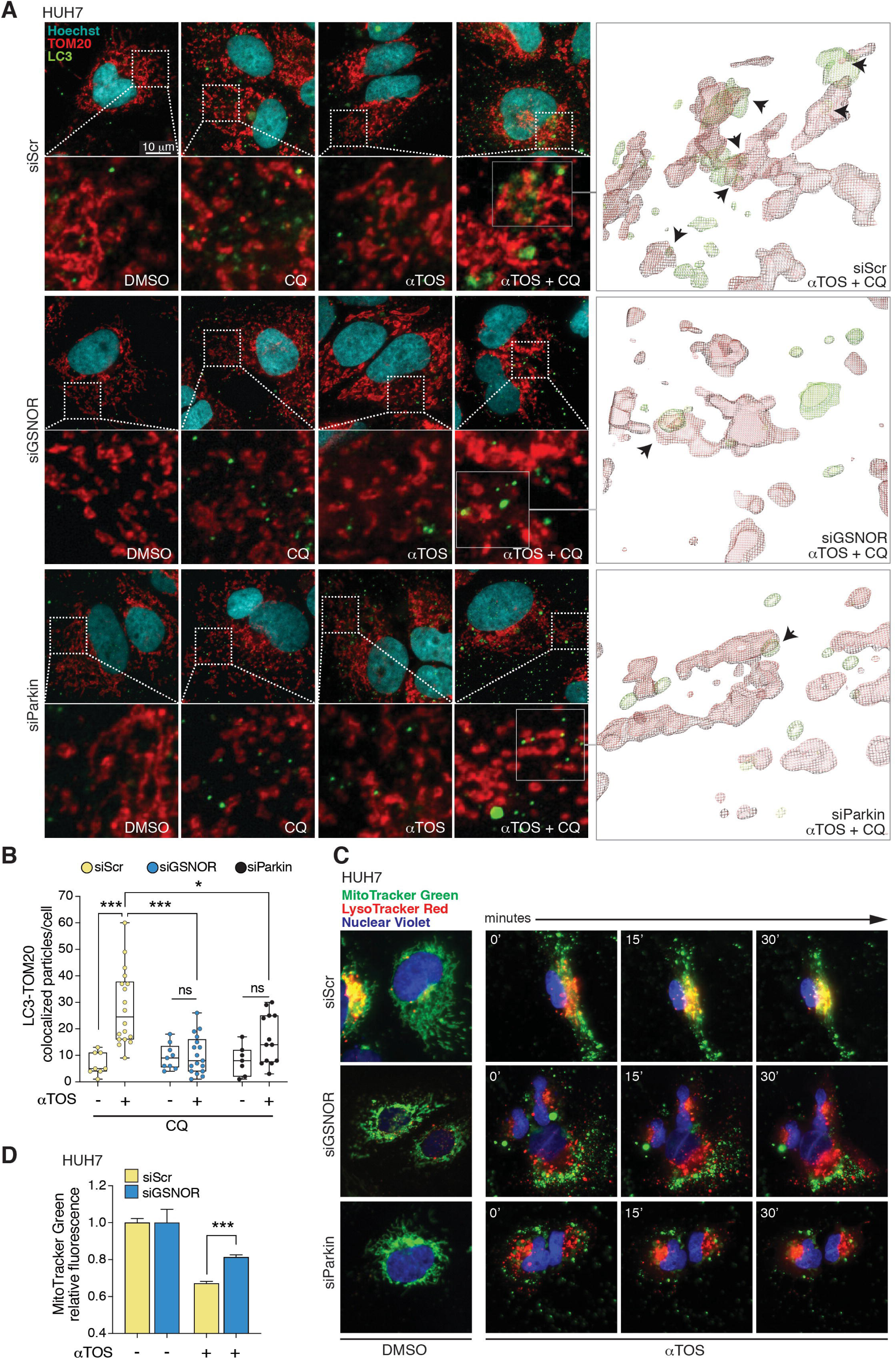
GSNOR-deficient hepatocellular carcinoma cells are insensitive to α TOS-induced mitophagy. **(A)** Mitophagy evaluation by fluorescence confocal microscopy of siScr, siGSNOR and siParkin HUH7 cells treated for 24 h with 40 μM αTOS and incubated with chloroquine (CQ) for 4 h to block mitochondrial degradation within autophagolysosomes. Mitochondria were labeled with an antibody against TOM20 (red), whereas autophagosomes were detected by using an anti-LC3 antibody (green). Hoechst 33342 (blue) was used to visualize nuclei. 3 central z-stacks (0.3 µm size) were merged in the microscopy pictures while >6 stacks were used for the 3D-rendering of TOM20 and LC3 signals (right panels). **(B)** Number of mitochondria colocalizing with LC3-positive puncta in CQ ± αTOS-treated cells calculated by Fiji analysis software using the open-source plugin ComDet v. 0.3.7. \**p* < 0.05; \*\*\**p* < 0.001; ns = not significant. At least 8 cells/experimental condition were counted. **(C)** Representative frames captured upon live-imaging fluorescence microscopy (Movies 7-12) of siScr and siGSNOR HUH7 stained with LysoTracker Red (red), MitoTracker Green (green) and Nuclear Violet (blue) to visualize lysosomes, mitochondria and nuclei, respectively. Cells, where indicated, were pre-treated for 24 h with 40 μM αTOS (or DMSO). After the staining, αTOS (or DMSO) was added before images acquisition. Images were acquired every 5 minutes for 90 minutes. **(D)** Mitochondrial mass analyzed by flow cytoflurometric detection of MitoTracker Green fluorescence of siScr and siGSNOR HUH7 cells treated for 24 h with 60 μM αTOS. Values represent the means ± SD of n = 3 independent experiments. \*\*\**p* < 0.001.

### 3.4. Mitophagy impairment contributes to αTOS cytotoxicity

We finally investigated the relationship between mitophagy and αTOS toxicity by analyzing cell death and survival upon GSNOR and/or Parkin downregulation (**Figure 5A**) in hepatocellular carcinoma cells. To this end, we treated HUH7 cells with αTOS for 24 h and evaluated cell death by combined staining with green-fluorescent calcein-AM (for living cells) and propidium iodide (for dead cells). Fluorescence microscopy analyses showed that GSNOR-deficient cells were more sensitive to αTOS cytotoxicity, confirming data previously obtained in our laboratory [39] (**Figure 5B, C**). Interestingly, we also observed that Parkin downregulation enhanced αTOS-induced cell death in siScr cell, whereas it did not produce any additional effect in siGSNOR cells, arguing for a *S*-nitrosylation and Parkin laying on the same pathways. We then repeated the same experiment in HepG2 cells (**Figure 5D**), confirming the same role of αTOS on cell viability in shGSNOR cells, where no further effects were observed in combination with siParkin (**Figure 5E**). This data suggested that, by sensitizing shScr cells to αTOS, Parkin silencing abolished the differences between GSNOR-deficient and proficient cells (**Figure 5E**). We finally confirmed these results by Cristal Violet survival assay of cells treated for 4 days with αTOS. GSNOR downregulation as well as Parkin silencing resulted in a lower survival rate if compared with GSNOR- or Parkin-proficient (shScr or siScr) counterparts (**Figure 5F**). In conclusion, these experiments reveal that it is possible to sensitize hepatocellular carcinoma cells to αTOS by inhibiting mitophagy. Moreover, our data suggest that *S*-nitrosylation contributes to cell susceptibility to αTOS by a mitophagy-dependent mechanism, inasmuch Parkin does not, as these cells are already mitophagy defective (**Figure 6**).

**Figure 5.**
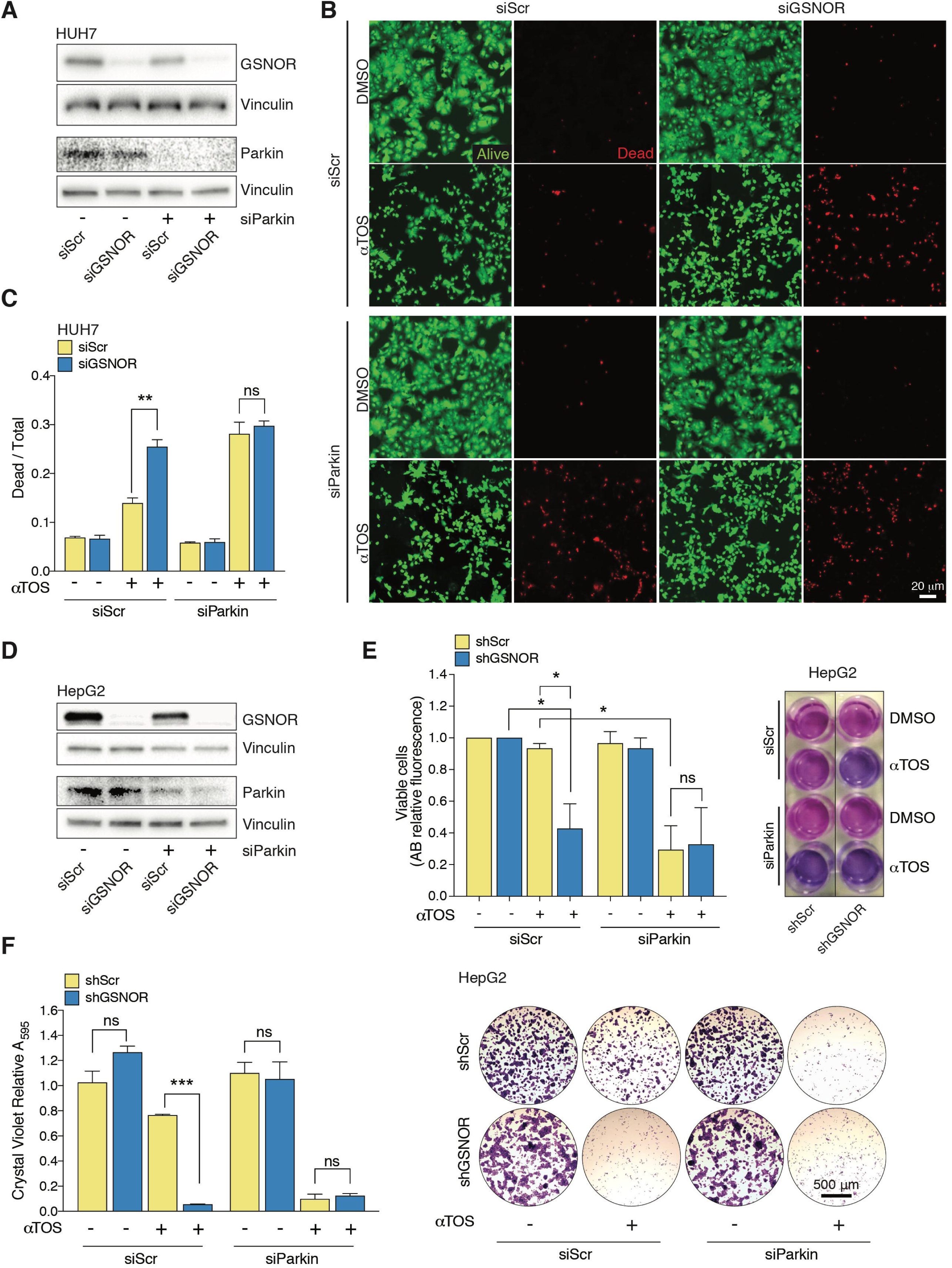
Mitophagy impairment contributes to GSNOR-deficient hepatocellular carcinoma sensitivity to αTOS. **(A)** Western blot analysis of GSNOR and Parkin in HUH7 cells transiently downregulating GSNOR (siGSNOR) and/or Parkin (siParkin). Vinculin was used as loading control. **(B)** Cell viability fluorescent assay performed in HUH7 cells singly or doubly transfected with siGSNOR or siParkin RNAs, and treated with 40 μM αTOS. Dead cells were stained with propidium iodide (red), whereas living cells were stained with calcein-AM (green). DMSO was selected as a control. **(C)** Cell death evaluation as ratio between dead and total cells and expressed as mean ± SEM of n = 5 different fields of n=3 independent experiments. \*\**p* < 0.01; ns = not significant. **(D)** Western blot analysis of GSNOR and Parkin in HepG2 cells singly or doubly transfected with siGSNOR or siParkin RNAs. Vinculin was used as loading control. **(E)** Analysis of cell viability upon staining with AlamarBlue in the same experimental setting as in **(D)**. DMSO was selected as a control. Viability is expressed as mean ± SEM of n = 3 independent experiments in triplicate. \**p* < 0.05; ns = not significant. **(F)** Cell survival analysis of shScr and shGSNOR HepG2 cells in in the same experimental setting as in **(D)**. Afterwards, cells were 5-fold diluted and maintained for 4 additional days in the presence or not of αTOS. Cells were fixed and stained with Crystal Violet at the end of the treatment. Representative pictures (right) and quantitative analysis (left) of Crystal Violet incorporation, performed after elution in methanol and measured at 595 nm. Data shown represent mean ± SD of n = 3 independent experiments. \*\*\**p* < 0.001. ns = not significant.

**Figure 6.**
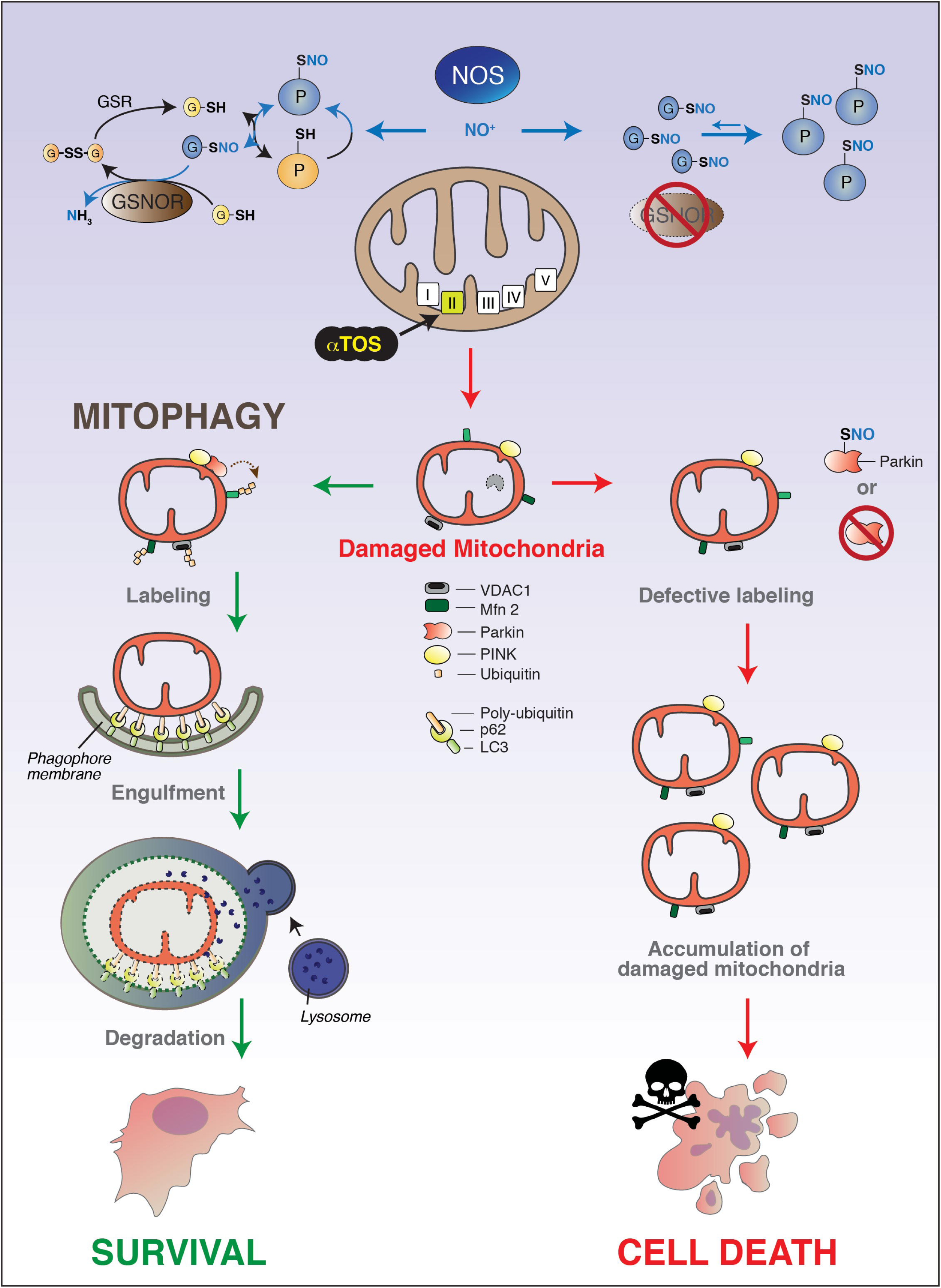
Working model of mitophagy-dependent α TOS toxicity in GSNOR-deficient cancer cells. Nitric Oxide (NO) is a signaling molecule physiologically produced by a class of nitric oxide synthases (NOS). The denitrosylase *S-*nitrosoglutathione reductase (GSNOR), by reducing the nitrosylated form of glutathione (GSH), *S-*nitrosoglutathione (GSNO), indirectly controls the extent of *S-*nitrosylated proteins (PSNO) (Upper left). GSNOR downregulation, a condition occurring during aging and in some cancer types, results in the accumulation of PSNOs (Upper right). αTOS, by targeting complex II of the mitochondrial electron transport chain, induces mitochondrial damage that, if not neutralized, could lead to cell death. In GSNOR-proficient cells, the selective recognition and degradation of mitochondria (mitophagy) is fully working and ensures a correct clearance of αTOS-damaged mitochondria (Center left). Damaged mitochondria are labeled by means of polyubiquitination of mitochondrial membrane-associated proteins (*e.g.* Mitofusin 2, Mfn2; Voltage-dependent anion channel, VDAC), carried out by E3 protein ubiquitin-ligases, among which Parkin represents one of the best characterized examples. Upon damage, PINK1 recruits Parkin onto mitochondria, this representing the starting point for ubiquitin-based labeling of damaged organelles. These are finally recognized by adaptor proteins (*e.g.* p62 and LC3); surrounded by a double membrane; engulfed, and degraded by fusing with a lysosome. Removal of defective mitochondria by mitophagy has pro-survival effects and contributes to chemoresistance (Bottom left). On the other hand, when GSNOR is mutated or loss, Parkin is inactivated by *S-*nitrosylation, becoming inactive and unable to be recruited in the proximity of damaged mitochondria, a condition similar to Parkin-deficient cells. In these conditions, αTOS treatment results in the accumulation of defective mitochondria, finally leading to cell death (Bottom right).

## 4. Discussion

Mechanisms underlying mitophagy are currently under intense investigation as they represent potential therapeutic targets due to the cytoprotective role of this process in cancer. In this regard, it has been demonstrated that the efficiency in clearing up drug-damaged mitochondria attenuates, or even abolishes, treatment effectiveness in several chemotherapy settings [17,20,48]. Starting from this evidence, many research groups developed the idea of using mitophagy inhibitors to enhance the efficacy of conventional chemotherapies [21]. Similarly, the identification of mitophagy-defective cancer subtypes advocates the application of mitochondrial-targeting therapies designed to selectively affect cancer cells survival, avoiding mitophagy-proficient cells [21]. Interestingly, the main characterized mitophagy regulators, *i.e.* the PINK1/Parkin system, BNIP3 and NIX, have been shown to be downregulated in several cancer types [16,25,49–52], suggesting that these could be targets of mitochondrial-directed therapies.

It is commonly known that Parkin-null mice develop liver cancer spontaneously [51]. Intriguingly, mice deficient for the denitrosylase GSNOR develop liver cancer as well [36,53], this pointing toward a common mechanism underlying tumorigenesis [54]. As a matter of fact, GSNOR-deficient cells show aberrant *S-*nitrosylation of Parkin, condition that compromises the E3-Ub ligase activity [34] and results in mitophagy defects [30]. Supported by the findings of this work, it is reasonable to include GSNOR-deficient hepatocellular carcinoma among the mitophagy-defective cancers.

In addition, we recently demonstrated that, due to mitochondrial electron transport chain rearrangements, GSNOR-deficient liver cancers cells are particularly susceptible to the mitochondrial complex II-targeting drug αTOS [39,41,55]. We here expanded the knowledge about the mechanism of action of αTOS, demonstrating for the first time that this toxin induces mitophagy, and that, similarly to other chemotherapics, the removal of damaged mitochondria through mitophagy dampens its toxicity. These discoveries are in line with previous data indicating lysosome instability at the bases of αTOS-induced apoptosis [56] in solid tumors, in particular colorectal [56] and breast [57,58] cancer. It is worth to note that breast cancer is actually characterized by GSNOR downregulation [38]. However, whether mitophagy defects contribute to αTOS efficacy in breast cancer eradication still needs to be elucidated. More generally, the effectiveness of mitocans in the treatment of mitophagy-defective tumors is still neglected, even though it might represent an intriguing field of investigation.

In conclusion, in the present study, we propose the use of αTOS as a promising tool toward GSNOR- and Parkin-defective, as well as other mitophagy-deficient tumors, and provide a molecular rationale to predict tumor response to mitocans.

## Acknowledgements

We would like to thank Laila Fisher for secretary assistance and Vanda Turcanova for the technical support. This work has been supported by Danish Cancer Society Grant Kræftens Bekæmpelses Videnskabelige Udvalg (KBVU) Grant [R146-A9414 and R231-A13855 to G. F, and R204-A12424 to D.D.Z.], Novo Nordisk Foundation (2018-0052550 to G.F), Associazione Italiana per la Ricerca sul Cancro (AIRC) Grant [IG20719] (to G.F.), LEO foundation (LF-OC-19-000004 to D.D.Z.), the Melanoma Research Alliance (to D.D.Z.). The laboratories in Copenhagen is part of the Center of Excellence in Autophagy, Recycling and Disease (CARD), funded by the Danish National Research Foundation (DNRF125).

## Author contributions

S.R. conceived and designed the study, performed the majority of experiments and interpreted the data; L.D.L. contributed to cell maintenance, treatments and performed survival assays; S.M. performed flow cytometry analyses; D.D.Z. and G.F. critically commented the manuscript and the figures; S.R. and G.F. wrote the manuscript.

## References

[1] J.Y. Chang, H.S. Yi, H.W. Kim, M. Shong, Dysregulation of mitophagy in carcinogenesis and tumor progression, Biochim. Biophys. Acta - Bioenerg. 1858 (2017) 633–640. doi: 10.1016/j.bbabio.2016.12.008.

[2] S. Vyas, E. Zaganjor, M.C. Haigis, Mitochondria and Cancer, Cell. 166 (2016) 555–566. doi: 10.1016/j.cell.2016.07.002.

[3] D.C. Chan, Mitochondria: Dynamic Organelles in Disease, Aging, and Development, Cell. 125 (2006) 1241–1252. doi: 10.1016/j.cell.2006.06.010.

[4] M. V Liberti, J.W. Locasale, The Warburg Effect: How Does it Benefit Cancer Cells?, Trends Biochem. Sci. 41 (2016) 211–218. doi: 10.1016/j.tibs.2015.12.001.

[5] J. Lu, M. Tan, Q. Cai, The Warburg effect in tumor progression: Mitochondrial oxidative metabolism as an anti-metastasis mechanism, Cancer Lett. 356 (2015) 156–164. doi: 10.1016/j.canlet.2014.04.001.

[6] M.G. Vander Heiden, L.C. Cantley, C.B. Thompson, Understanding the Warburg effect: the metabolic requirements of cell proliferation., Science. 324 (2009) 1029–33. doi: 10.1126/science.1160809.

[7] L. Tochhawng, S. Deng, S. Pervaiz, C.T. Yap, Redox regulation of cancer cell migration and invasion, Mitochondrion. 13 (2013) 246–253. doi: 10.1016/j.mito.2012.08.002.

[8] M.C. De Santis, P.E. Porporato, M. Martini, A. Morandi, Signaling Pathways Regulating Redox Balance in Cancer Metabolism, Front. Oncol. 8 (2018) 126. doi: 10.3389/fonc.2018.00126.

[9] A. Rasola, L. Neckers, D. Picard, Mitochondrial oxidative phosphorylation TRAP(1)ped in tumor cells, Trends Cell Biol. 24 (2014) 455–463. doi: 10.1016/j.tcb.2014.03.005.

[10] C.R. Reczek, N.S. Chandel, ROS Promotes Cancer Cell Survival through Calcium Signaling, Cancer Cell. 33 (2018) 949–951. doi: 10.1016/j.ccell.2018.05.010.

[11] D.R. Green, L. Galluzzi, G. Kroemer, Metabolic control of cell death, Science (80-.). 345 (2014) 1250256–1250256. doi: 10.1126/science.1250256.

[12] A. Kasahara, L. Scorrano, Mitochondria: from cell death executioners to regulators of cell differentiation, Trends Cell Biol. 24 (2014) 761–770. doi: 10.1016/j.tcb.2014.08.005.

[13] R.J. De Berardinis, N.S. Chandel, Fundamentals of cancer metabolism, Sci. Adv. 2 (2016) 107–111. doi: 10.1126/sciadv.1600200.

[14] T. Han, D. Kang, D. Ji, X. Wang, W. Zhan, M. Fu, H.B. Xin, J. Bin Wang, How does cancer cell metabolism affect tumor migration and invasion?, Cell Adhes. Migr. 7 (2013) 395–403. doi: 10.4161/cam.26345.

[15] J.Y. Chang, H.-S. Yi, H.-W. Kim, M. Shong, Dysregulation of mitophagy in carcinogenesis and tumor progression, Biochim. Biophys. Acta - Bioenerg. 1858 (2017) 633–640. doi: 10.1016/j.bbabio.2016.12.008.

[16] E. Villa, E. Proïcs, C. Rubio-Patiño, S. Obba, B. Zunino, J.P. Bossowski, R.M. Rozier, J. Chiche, L. Mondragón, J.S. Riley, S. Marchetti, E. Verhoeyen, S.W.G. Tait, J.E. Ricci, Parkin-Independent Mitophagy Controls Chemotherapeutic Response in Cancer Cells, Cell Rep. 20 (2017) 2846–2859. doi: 10.1016/j.celrep.2017.08.087.

[17] L. Hardy, M. Frison, M. Campanella, Breast cancer cells exploit mitophagy to exert therapy resistance, Oncotarget. 9 (2018) 14040–14041. doi: 10.18632/oncotarget.24533.

[18] M. Vara-Perez, B. Felipe-Abrio, P. Agostinis, Mitophagy in Cancer: A Tale of Adaptation, Cells. 8 (2019) 493. doi: 10.3390/cells8050493.

[19] C. Yan, L. Luo, C.-Y. Guo, S. Goto, Y. Urata, J.-H. Shao, T.-S. Li, Doxorubicin-induced mitophagy contributes to drug resistance in cancer stem cells from HCT8 human colorectal cancer cells, Cancer Lett. 388 (2017) 34–42. doi: 10.1016/j.canlet.2016.11.018.

[20] P. Bhat, J. Kriel, B. Shubha Priya, Basappa, N.S. Shivananju, B. Loos, Modulating autophagy in cancer therapy: Advancements and challenges for cancer cell death sensitization, Biochem. Pharmacol. 147 (2018) 170–182. doi: 10.1016/j.bcp.2017.11.021.

[21] H.Y. Chiu, E.X.Y. Tay, D.S.T. Ong, R. Taneja, Mitochondrial dysfunction at the centre of cancer therapy, Antioxid. Redox Signal. 00 (2019) 1–22. doi: 10.1089/ars.2019.7898.

[22] C. Zhao, R. He, M. Shen, F. Zhu, M. Wang, Y. Liu, H. Chen, X. Li, R. Qin, PINK1/Parkin-Mediated Mitophagy Regulation by Reactive Oxygen Species Alleviates Rocaglamide A-Induced Apoptosis in Pancreatic Cancer Cells, Front. Pharmacol. 10 (2019) 1–13. doi: 10.3389/fphar.2019.00968.

[23] L. Song, Y. Huang, X. Hou, Y. Yang, S. Kala, Z. Qiu, R. Zhang, L. Sun, PINK1/Parkin-Mediated Mitophagy Promotes Resistance to Sonodynamic Therapy, Cell. Physiol. Biochem. 49 (2018) 1825–1839. doi: 10.1159/000493629.

[24] J. Okami, D.M. Simeone, C.D. Logsdon, Silencing of the Hypoxia-Inducible Cell Death Protein BNIP3 in Pancreatic Cancer, Cancer Res. 64 (2004) 5338 LP – 5346. doi: 10.1158/0008-5472.CAN-04-0089.

[25] A.H. Chourasia, K.F. Macleod, Tumor suppressor functions of BNIP3 and mitophagy, Autophagy. 11 (2015) 1937–1938. doi: 10.1080/15548627.2015.1085136.

[26] T. Kimura, Y. Takabatake, A. Takahashi, Y. Isaka, Chloroquine in Cancer Therapy: A Double-Edged Sword of Autophagy, Cancer Res. 73 (2013) 3 LP – 7. doi: 10.1158/0008-5472.CAN-12-2464.

[27] J. Zhou, G. Li, Y. Zheng, H.M. Shen, X. Hu, Q.L. Ming, C. Huang, P. Li, N. Gao, A novel autophagy/mitophagy inhibitor liensinine sensitizes breast cancer cells to chemotherapy through DNM1L-mediated mitochondrial fission, Autophagy. 11 (2015) 1259–1279. doi: 10.1080/15548627.2015.1056970.

[28] A. Managò, L. Leanza, L. Carraretto, N. Sassi, S. Grancara, R. Quintana-Cabrera, V. Trimarco, A. Toninello, L. Scorrano, L. Trentin, G. Semenzato, E. Gulbins, M. Zoratti, I. Szabò, Early effects of the antineoplastic agent salinomycin on mitochondrial function, Cell Death Dis. 6 (2015). doi: 10.1038/cddis.2015.263.

[29] C. Yan, T.S. Li, Dual role of mitophagy in cancer drug resistance, Anticancer Res. 38 (2018) 617–621. doi: 10.21873/anticanres.12266.

[30] S. Rizza, S. Cardaci, C. Montagna, G. Di Giacomo, D. De Zio, M. Bordi, E. Maiani, S. Campello, A. Borreca, A.A. Puca, J.S. Stamler, F. Cecconi, G. Filomeni, S -nitrosylation drives cell senescence and aging in mammals by controlling mitochondrial dynamics and mitophagy, Proc. Natl. Acad. Sci. U. S. A. 10 (2018) E3388–E3397. doi: 10.1007/s10986-018-9401-8.

[31] S. Rizza, G. Filomeni, Denitrosylate and live longer: how ADH5/GSNOR links mitophagy to aging, Autophagy. 14 (2018) 1285–1287. doi: 10.1080/15548627.2018.1475818.

[32] D.T. Hess, A. Matsumoto, S.-O.O. Kim, H.E. Marshall, J.S. Stamler, Protein S-nitrosylation: purview and parameters, Nat.Rev.Mol.Cell Biol. 6 (2005) 150–166. doi: 10.1038/nrm1569.

[33] D.T. Hess, J.S. Stamler, Regulation by S-nitrosylation of protein post-translational modification, J. Biol. Chem. 287 (2012) 4411–4418. doi: 10.1074/jbc.R111.285742.

[34] K.K.K. Chung, B. Thomas, X. Li, O. Pletnikova, J.C. Troncoso, L. Marsh, V.L. Dawson, T.M. Dawson, S-nitrosylation of parkin regulates ubiquitination and compromises parkin’s protective function., Science (80-.). 304 (2004) 1328–31. doi: 10.1126/science.1093891.

[35] T. Nakamura, S.A. Lipton, S-Nitrosylation of Critical Protein Thiols Mediates Protein Misfolding and Mitochondrial Dysfunction in Neurodegenerative Diseases, Antioxid. Redox Signal. 14 (2010) 1479–1492. doi: 10.1089/ars.2010.3570.

[36] W. Wei, B. Li, M.A. Hanes, S. Kakar, X. Chen, L. Liu, S-nitrosylation from GSNOR deficiency impairs DNA repair and promotes hepatocarcinogenesis., Sci. Transl. Med. 2 (2010) 19ra13. doi: 10.1126/scitranslmed.3000328.

[37] C.H. Tang, W. Wei, M.A. Hanes, L. Liu, Hepatocarcinogenesis driven by GSNOR deficiency is prevented by iNOS inhibition, Cancer Res. 73 (2013) 2897–2904. doi: 10.1158/0008-5472.CAN-12-3980.

[38] A. Cañas, L.M. López-Sánchez, J. Peñarando, A. Valverde, F. Conde, V. Hernández, E. Fuentes, C. López-Pedrera, J.R. de la Haba-Rodríguez, E. Aranda, A. Rodríguez-Ariza, Altered S-nitrosothiol homeostasis provides a survival advantage to breast cancer cells in HER2 tumors and reduces their sensitivity to trastuzumab, Biochim. Biophys. Acta - Mol. Basis Dis. 1862 (2016) 601–610. doi: 10.1016/j.bbadis.2016.02.005.

[39] S. Rizza, C. Montagna, S. Cardaci, E. Maiani, G. Di Giacomo, V. Sanchez-Quiles, B. Blagoev, A. Rasola, D. De Zio, J.S. Stamler, F. Cecconi, G. Filomeni, S-nitrosylation of the mitochondrial chaperone TRAP1 sensitizes hepatocellular carcinoma cells to inhibitors of succinate dehydrogenase, Cancer Res. 76 (2016) 4170–4182. doi: 10.1158/0008-5472.CAN-15-2637.

[40] J. Neuzil, L.-F. Dong, J. Rohlena, J. Truksa, S.J. Ralph, Classification of mitocans, anti-cancer drugs acting on mitochondria, Mitochondrion. 13 (2013) 199–208. doi: 10.1016/j.mito.2012.07.112.

[41] L.-F. Dong, P. Low, J.C. Dyason, X.-F. Wang, L. Prochazka, P.K. Witting, R. Freeman, E. Swettenham, K. Valis, J. Liu, R. Zobalova, J. Turanek, D.R. Spitz, F.E. Domann, I.E. Scheffler, S.J. Ralph, J. Neuzil, Alpha-tocopheryl succinate induces apoptosis by targeting ubiquinone-binding sites in mitochondrial respiratory complex II., Oncogene. 27 (2008) 4324–35. doi: 10.1038/onc.2008.69.

[42] J. Neuzil, Vitamin E succinate and cancer treatment: a vitamin E prototype for selective antitumour activity., Br. J. Cancer. 89 (2003) 1822–6. doi: 10.1038/sj.bjc.6601360.

[43] J. Neuzil, T. Weber, N. Gellert, C. Weber, Selective cancer cell killing by alpha-tocopheryl succinate., Br. J. Cancer. 84 (2001) 87–89. doi: 10.1054/bjoc.2000.1559.

[44] J. Schindelin, I. Arganda-Carreras, E. Frise, V. Kaynig, M. Longair, T. Pietzsch, S. Preibisch, C. Rueden, S. Saalfeld, B. Schmid, J.Y. Tinevez, D.J. White, V. Hartenstein, K. Eliceiri, P. Tomancak, A. Cardona, Fiji: An open-source platform for biological-image analysis, Nat. Methods. 9 (2012) 676–682. doi: 10.1038/nmeth.2019.

[45] E.F. Pettersen, T.D. Goddard, C.C. Huang, G.S. Couch, D.M. Greenblatt, E.C. Meng, T.E. Ferrin, UCSF Chimera—A visualization system for exploratory research and analysis, J. Comput. Chem. 25 (2004) 1605–1612. doi: 10.1002/jcc.20084.

[46] O.H. Lowry, N.J. Rosebrough, A.L. Farr, R.J. Randall, PROTEIN MEASUREMENT WITH THE FOLIN PHENOL REAGENT, J. Biol. Chem.. 193 (1951) 265–275. http://www.jbc.org/content/193/1/265.short.

[47] M. Mauthe, I. Orhon, C. Rocchi, X. Zhou, M. Luhr, K.-J. Hijlkema, R.P. Coppes, N. Engedal, M. Mari, F. Reggiori, Chloroquine inhibits autophagic flux by decreasing autophagosome-lysosome fusion, Autophagy. 14 (2018) 1435–1455. doi: 10.1080/15548627.2018.1474314.

[48] S.-F. Wang, M.-S. Chen, Y.-C. Chou, Y.-F. Ueng, P.-H. Yin, T.-S. Yeh, H.-C. Lee, Mitochondrial dysfunction enhances cisplatin resistance in human gastric cancer cells via the ROS-activated GCN2-eIF2α-ATF4-xCT pathway, Oncotarget. 7 (2016). doi: 10.18632/oncotarget.12356.

[49] E. Villa, E. Proïcs, C. Rubio-Patiño, S. Obba, B. Zunino, J.P. Bossowski, R.M. Rozier, J. Chiche, L. Mondragón, J.S. Riley, S. Marchetti, E. Verhoeyen, S.W.G. Tait, J.-E. Ricci, Parkin-Independent Mitophagy Controls Chemotherapeutic Response in Cancer Cells, Cell Rep. 20 (2017) 2846–2859. doi: 10.1016/j.celrep.2017.08.087.

[50] N. Yao, C. Wang, N. Hu, Y. Li, M. Liu, Y. Lei, M. Chen, L. Chen, C. Chen, P. Lan, W. Chen, Z. Chen, D. Fu, W. Ye, D. Zhang, Inhibition of PINK1/Parkin-dependent mitophagy sensitizes multidrug-resistant cancer cells to B5G1, a new betulinic acid analog, Cell Death Dis. 10 (2019) 232. doi: 10.1038/s41419-019-1470-z.

[51] J.P. Bernardini, M. Lazarou, G. Dewson, Parkin and mitophagy in cancer, Oncogene. 36 (2016) 1315. doi: 10.1038/onc.2016.302.

[52] J. Zhang, P.A. Ney, Role of BNIP3 and NIX in cell death, autophagy, and mitophagy, Cell Death Differ. 16 (2009) 939. doi: 0.1038/cdd.2009.16.

[53] W. Wei, Z. Yang, C.H. Tang, L. Liu, Targeted deletion of GSNOR in hepatocytes of mice causes nitrosative inactivation of O6-alkylguanine-dna alkyltransferase and increased sensitivity to genotoxic diethylnitrosamine, Carcinogenesis. 32 (2011) 973–977. doi: 10.1093/carcin/bgr041.

[54] S. Rizza, G. Filomeni, Tumor Suppressor Roles of the Denitrosylase GSNOR, Crit. Rev. Oncog. 21 (2016) 433–445. doi: 10.1615/CritRevOncog.2017021074.

[55] L.F. Dong, V.J.A. Jameson, D. Tilly, J. Cerny, E. Mahdavian, A. Marín-Hernández, L. Hernández-Esquivel, S. Rodríguez-Enríquez, J. Stursa, P.K. Witting, B. Stantic, J. Rohlena, J. Truksa, K. Kluckova, J.C. Dyason, M. Ledvina, B.A. Salvatore, R. Moreno-Sánchez, M.J. Coster, S.J. Ralph, R.A.J. Smith, J. Neuzil, Mitochondrial targeting of vitamin E succinate enhances its pro-apoptotic and anti-cancer activity via mitochondrial complex II, J. Biol. Chem. 286 (2011) 3717–3728. doi: 10.1074/jbc.M110.186643.

[56] J. Neuzil, M. Zhao, G. Ostermann, M. Sticha, N. Gellert, C. Weber, J.W. Eaton, U.T. Brunk, Alpha-tocopheryl succinate, an agent with in vivo anti-tumour activity, induces apoptosis by causing lysosomal instability, Biochem. J. 362 (2002) 709–715. doi: 10.1042/0264-6021:3620709.

[57] M.P. Malafa, L.T. Neitzel, Vitamin E Succinate Promotes Breast Cancer Tumor Dormancy, J. Surg. Res. 93 (2000) 163–170. doi: 10.1006/jsre.2000.5948.

[58] L.F. Dong, R. Freeman, J. Liu, R. Zobaiova, A. Marin-Hernandez, M. Stantic, J. Rohlena, K. Valis, S. Rodriguez-Enriquez, B. Butcher, J. Goodwin, U.T. Brunk, P.K. Witting, R. Moreno-Sanchez, I.E. Scheffler, S.J. Raiph, J. Neuzil, Suppression of tumor growth in vivo by the mitocan α-tocopheryl succinate requires respiratory complex II, Clin. Cancer Res. 15 (2009) 1593–1600. doi: 10.1158/1078-0432.CCR-08-2439.

